# Semblance of heterogeneity in collective cell migration

**DOI:** 10.1101/122374

**Authors:** Linus J. Schumacher, Philip K. Maini, Ruth E. Baker

## Abstract

Cell population heterogeneity is increasingly a focus of inquiry in biological research. For example, cell migration studies have investigated the heterogeneity of invasiveness and taxis in development, wound healing, and cancer. However, relatively little effort has been devoted to explore when heterogeneity is mechanistically relevant and how to reliably measure it. Statistical methods from the animal movement literature offer the potential to analyse heterogeneity in collections of cell tracking data. A popular measure of heterogeneity, which we use here as an example, is the distribution of delays in directional cross-correlation. Employing a suitably generic, yet minimal, model of collective cell movement in three dimensions, we show how using such measures to quantify heterogeneity in tracking data can result in the inference of heterogeneity where there is none. Our study highlights a potential pitfall in the statistical analysis of cell population heterogeneity, and we argue this can be mitigated by the appropriate choice of null models.

**Highlights:**

- groups of identical cells appear heterogeneous due to limited sampling and experimental repeatability
- heterogeneity bias increases with attraction/repulsion between cells
- movement in confined environments decreases apparent heterogeneity
- hypothetical applications in neural crest and *in vitro* cancer systems

**In Brief:** We use a mathematical model to show how cell populations can appear heterogeneous in their migratory characteristics, even though they are made up of identically-behaving individual cells. This has important consequences for the study of collective cell migration in areas such as embryo development or cancer invasion.

## 1 Introduction

Collective migration of cell populations plays an important role in development, regeneration, and disease. Evidence is mounting that population heterogeneity functionally contributes to the collective behaviour of cells in many systems. One form of heterogeneity that is frequently studied is that of leader and follower cell states within a population. This has been investigated in the migration of the zebrafish lateral line primordium (Streichan *et al.*, 2011), Drosophila border cells (Inaki *et al.*, 2012), neural crest cells (McLennan *et al.*, 2012, 2015a), as well as neutrophils and T-cells (Lim *et al.*, 2015), to mention only a few examples.

However, leader-follower heterogeneity is not found in all collectively migrating cell populations, and mathematical models are able to produce collective migration of a group of identical agents, calling into question the need for such heterogeneity (see discussion in Box 2, Schumacher *et al.*, 2016). In addition, evidence for the plasticity of leader cell states in neural crest (McLennan *et al.*, 2012, 2015b), as well as Drosophila border cell migration (Rørth, 2012), indicates that population heterogeneity may often emerge from the interaction of cells with microenvironmental signals, and therefore possesses a degree of plasticity. Recent theoretical work has also shown that (proliferative) heterogeneity can arise from constraints, such as confinement, alone (Smadbeck and Stumpf, 2016). Together these pieces of evidence raise the question of how cell interactions and microenvironmental conditions can induce, promote, or otherwise affect, the heterogeneity of a cell population, and our ability to measure it.

Cell tracking data provide a major source for evidence of heterogeneity of movement. Advances in threedimensional (3D) imaging and computational tracking of complete cell populations *in vivo* or in realistic *in vitro* assays give rise to rich datasets amenable to measuring distributions of statistics of interest. For example, recent efforts by Sharma et *al*. (2015) have drawn on methods from the literature on animal movement (Nagy *et al.*, 2010) to measure cross-correlational delay times of mammalian cell cohorts moving in a 3D extracellular matrix gel. Such methods could be of great interest, for example to characterise the invasiveness of cancer cells from different tumor samples and assess the efficacy of potential metastasis-inhibiting treatments. However, unlike the animal experiments for which such methods were initially developed, cell biology assays are limited in observation time, and typically cannot be repeated with the same cells while retaining individual identification. Both these shortcomings result in smaller datasets, which are more prone to spurious correlations from chance.

As researchers we are faced with the problem of when to include heterogeneity in our mechanistic descriptions of collective cell migration, and when not. In short, when we observe what appears to be heterogeneity, is this an intrinsic property of the system, an emergent phenomenon, or a statistical artefact? Here, we take inspiration from previous studies to highlight potential pitfalls in the analysis of tracking data from cell collectives. We develop a suitably versatile model for 3D collective cell migration and use this to generate trajectory data which we, in turn, analyse with a commonly used measure of heterogeneity, the distribution of delay times in directional cross-correlation. By showing that, under this protocol, we can measure heterogeneity in the form of leader-follower relationships despite individuals being identical, we demonstrate that care must be taken when using this or similar measures of heterogeneity. Our results highlight that appropriate null models must be carefully dedicated to particular experiments to assess the statistical significance of observed correlations. By analysing the systematic bias of our analysis toward apparent heterogeneity as a function of model parameters and boundary conditions, we propose two hypothetical applications of our work that could be tested using in vivo and *in vitro* cell tracking data. We conclude by discussing alternative measures of heterogeneity, further model extensions, and potential avenues for the development of statistical tests.

## 2. Results

### A generic model for collective cell migration

Grégoire *et al.* (2003) have previously developed a self-propelled particle (SPP) model for collective migration in two dimensions. We here extend this model to three dimensions, but for simplicity first explain the basic model components in two dimensions. The direction of movement, *θ*, of cell *i* at time *t* + 1 is given by

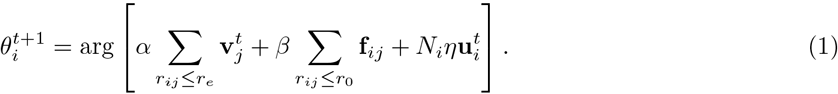

In the above equation, three terms independently influence the direction of movement, variously depending on the separation, *r_ij_*, between cells *i* and *j*: (1) alignment with the movement direction of all neighboring cells (within distance *r_e_*, where 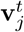 is the velocity of cell *j* at time *t*, for simplicity assumed to be of fixed speed), scaled by parameter *α* ≥ 0; (2) intercellular forces, **f***_ij_*, i.e., attraction/repulsion toward/away from neighboring cells (within distance *r*_0_), scaled by parameter *β* ≥ 0; (3) noise, here chosen to be a random vector on the unit sphere, 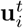, which scales with the number of neighbors, *N_i_* (within distance *r*_0_, including cell *i* itself), to represent the uncertainty in the forces from neighboring cells (Grégoire *et al.*, 2003). The noise term is controlled by parameter *η* ≥ 0, and without loss of generality we have set *η* = 1 in the simulations presented here. Interactions between cells are further restricted to nearest Voronoi neighbors only, even if more cells are within the respective distance cut-off. Fig. S1 illustrates the various interaction zones.

### Extension to three dimensions

In the 3D model we have developed here, the direction of movement is parameterised by two angles, the azimuthal angle, *θ* (from −*π* to *π*), and the polar angle, *ϕ* (from 0 to *π*), which are updated according to

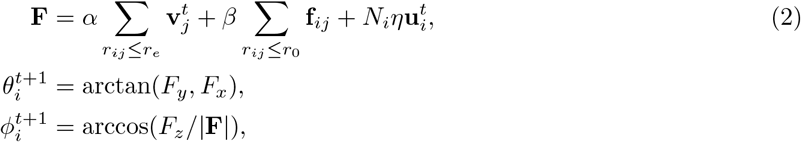

where the interactions in (2) are again between nearest neighbors only. Cell positions for cell *i* are updated via 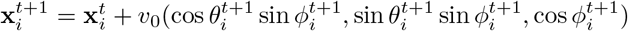, with fixed speed *υ*_0_ = 0.05, respecting boundary conditions. For details on the form of the intercellular forces see Intercellular forces. In this paper, we have implemented both free (cells are unconfined) and no-flux (reflective) boundary conditions. A similar 3D model has also been developed by Sharma *et al.* (2015), with small but important differences in the model implementation. Crucially, we have kept the original idea by Grégoire *et al.* (2003) to restrict interactions to nearest Voronoi neighbors.

### Intrinsic heterogeneity

In the work shown here, we primarily focus on homogeneous populations and how they can appear heterogeneous. To explicitly include heterogeneity, we repeated a subset of our simulations with a few “informed” cells, or cells in a leader state. The randomly chosen subset of informed cells align not with their neighbours, but with a prescribed direction (here chosen to be the *x*-axis without loss of generality). This preferred direction could represent, for example, a chemoattractant gradient or other directional cue. The alignment strength, *α*, is the same as for the other cells, and other cells still align with the informed cell if they are neighbouring them. We chose 10% of cells to be in such a leader state, as similarly small fractions have been reported sufficient to affect the overall population behaviour in migrating cell populations (McLennan *et al.*, 2015a) and active systems more generally (Yllanes and Marchetti, 2017).

### Computational experiments

The mathematical model has been implemented in Matlab. Unless stated otherwise, 3D simulations were run without confinement (free boundary conditions), for 100 cells whose initial positions were chosen randomly from within a cube of edge length *L* = 2. To alleviate the influence of transient behaviour dependent on initial conditions, we run simulations for 1000 time-steps and ignore the first 500 time-steps in our analysis, inspecting the order parameter (see Global model behaviour) plotted against time to validate that this is sufficient to reach a (quasi) steady-state. We find that groups of cells move in an ordered (globally aligned) manner for high values of *α* and *β* (Fig. S3). Example trajectories (Fig. 1) reveal moving streams and slowly translating clusters that break apart as the strength of intercellular forces, *β*, becomes comparable to the noise strength, *η* = 1, or lower. At low *α* and high *β*, groups of cells are in static, positionally ordered arrangements (Fig. 1).

**Figure 1:**
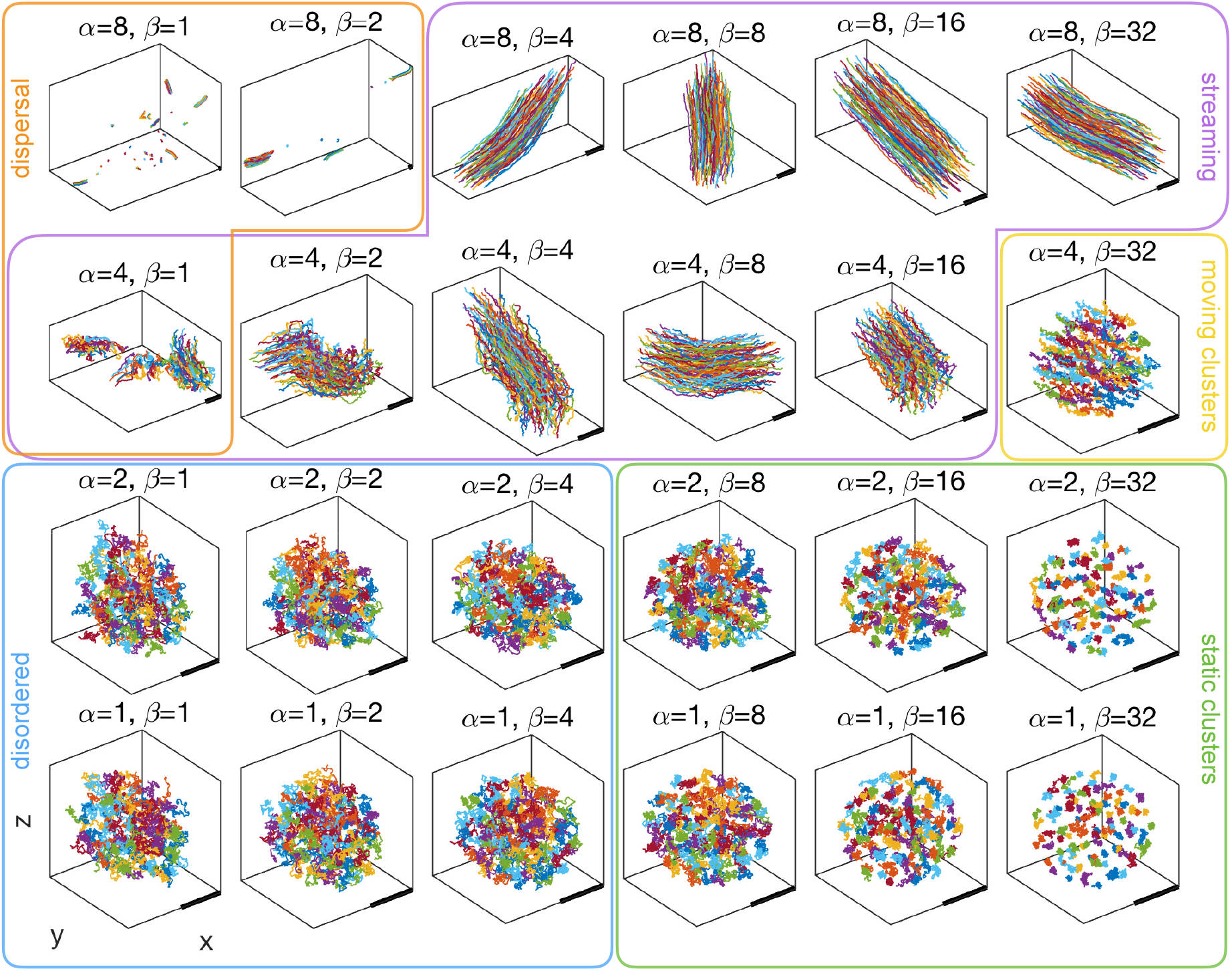
Example trajectories of simulations with *N* = 100 cells and free boundary conditions, showing a range of migratory behaviours achievable with our generic model, such as dispersal, streaming, moving, and static clusters, as well as disordered arrangements (labeled by visual inspection). Different model behaviours are achieved by varying the alignment strength, *α*, and the strength of attraction/repulsion, *β*. See also Eqn. (2). Here, 100 time-steps are shown after simulations have run for 500 time-steps, starting from random initial positions (see main text for details). Scale bar shows *L* = 1.

### Delay correlation analysis

To measure heterogeneity of movement within a group of moving cells, we measure the extent to which cells are following one another using delay correlation analysis. For two cells *i* and *j* the directional cross-correlation at lag *τ* calculated as

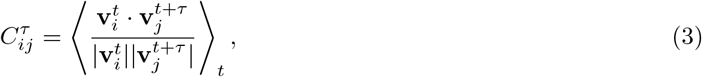

where · denotes the scalar product and 〈…〉*_t_* time-averaging. The peak-delay time, *τ_C_*, for a pair of cells is that for which 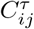 is highest in the observed data. By binning the peak-delay times for all cells in the population, we obtain the sample distribution *P*(*τ_C_*), so that *P* (*τ*_1_) = 1 would mean that all pairs of cells have their highest directional cross-correlation at lag *τ*_1_, and *P*(*τ*_0_) = 0 means no pair of cells has peak correlation at lag *τ*_0_ (note that *P*(*τ_C_*) is symmetric about *τ* = 0 since 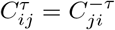). Thus, *P* (*τ_C_*) constitutes a population-scale measure of heterogeneity. To compare heterogeneity between multiple populations, we further summarise the width of the distribution by its standard deviation, *σ*(*τ_C_*).

To only consider cohesively moving populations, we restrict our analysis to model realisations with directional order above a minimum threshold, which we set at Φ_min_ = 0.1. Within those simulation results, we then find peak-delay times for any pairs of cells which have been within interaction radius *r*_0_ and have directional cross-correlation of *C_min_* = 0.5 or greater at at least one time-point. An illustrative example of the resulting distribution of peak-delay times is shown in Fig. 2(A), indicating that the width of the distribution increases with *β*. To test whether some cells showed consistently non-zero lag times (either leading ahead or following behind others), we also calculate the distribution of peak-delay times for the directional correlation between individual cells and all other cells in the population. An example of the individual distributions is shown in Fig. 2(B). For a more comprehensive view, we calculate the standard deviation of the peak-delay distribution, *σ*(*τ_C_*), as a measure of the width of the distribution, which can be seen to increase consistently with increasing *β* over a range of values for *α* (Fig. 2(C)). The standard deviation is also high for low values of *β* at some values of *α*, but this trend is not consistent across the range of *α* values. Increasing the alignment strength, *α*, for fixed *β* does not increase the measure of heterogeneity. In summary, stronger attraction/repulsion between cells leads to increased apparent heterogeneity.

**Figure 2:**
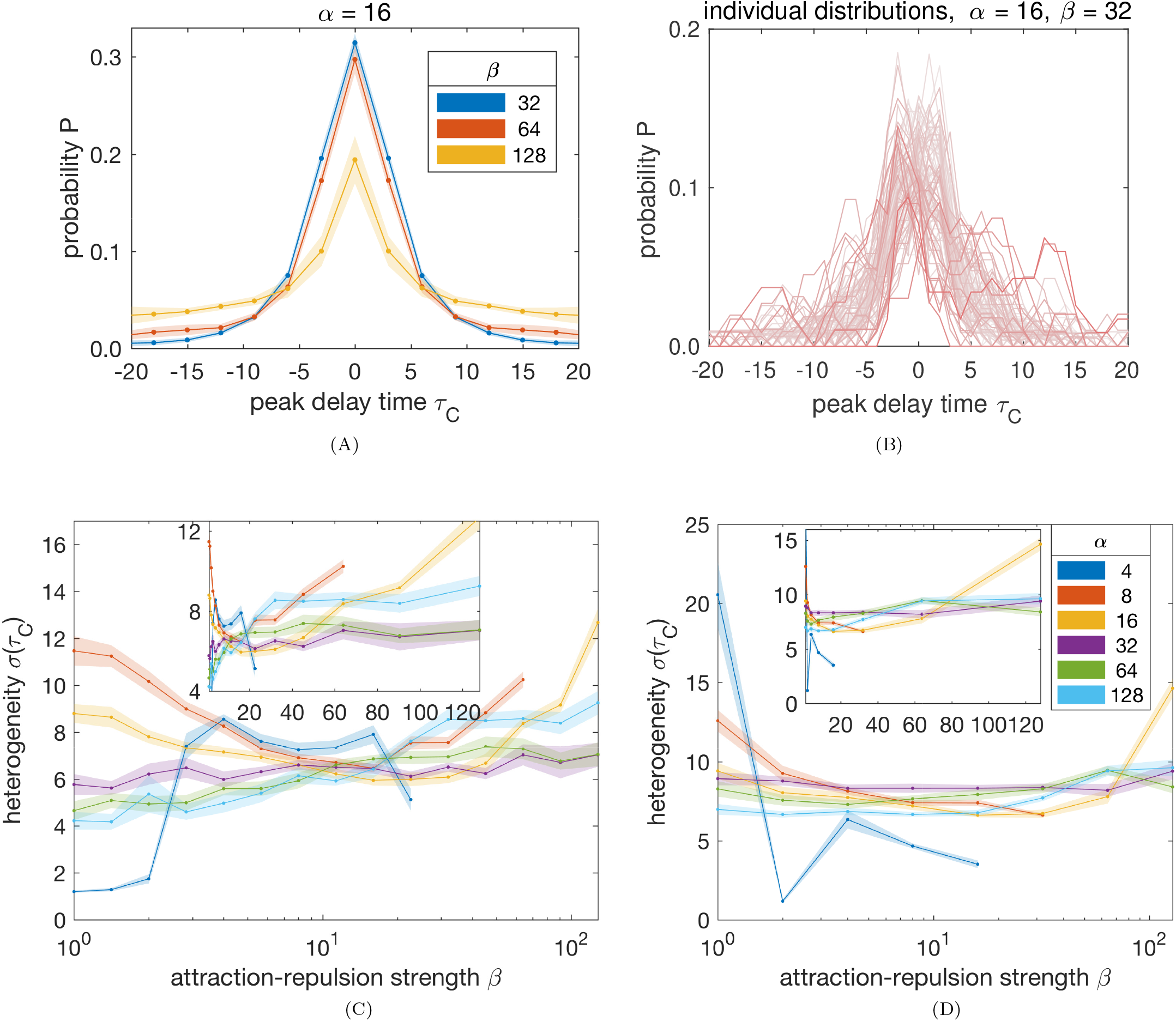
(A) Peak-delay distributions of model simulations with *α* = 16, for varying values of *β*. Points show the average for each bin over 10 simulations, shaded area shows the standard deviation. Peaks in directional cross-correlation between cells were calculated over 500 time-steps after 500 time-steps burn-in. (B) Peak-delay distributions of individual cells (smoothed) for one simulation of the chosen parameter combination. Distributions with higher absolute mean are shown in red, distributions with lower absolute mean (closer to zero) are shown in lighter shades of red. (C-D) Standard deviation of peak-delay distributions, *σ*(*τ_C_*), plotted against strength of intercellular forces, *β*, for varying values of alignment strength, *α*, calculated from simulations of homogenous populations (C), and from simulations including heterogeneity in the form of informed cells (D). Points show mean of 10 simulations, shaded areas show standard error of the means. The horizontal axes on the main plots show log-scale in *β*, while the insets show a linear scale, highlighting the systematic increase in heterogeneity over a wide range of β (more pronounced in C). Any missing data were excluded due to simulations not being ordered or sufficiently correlated (see main text for details).

When we included heterogeneity in our simulations, we found that the increase in apparent heterogeneity is less strong for high alignment strength (Fig. 2D). This can be understood as strong alignment with a prescribed direction suppressing directional fluctuations. The model with heterogeneity fit the experimental data (see Biological applications and Comparison with experimental data) at lower values of the interaction strength parameter (Fig. B1) than the homogenous model. This illustrates the cohesive effect that even small numbers of cells in a leader state can have on the population.

##### Biological applications

###### Neural crest migration

Neural crest (NC) cells display a wide range of migratory patterns in different organisms and embryonal locations, yet a unified mechanistic understanding has so far eluded the research community. For example, in chick cranial NC migration, data on cell behaviour and gene expression, as well as mathematical modelling, have suggested a collective migration mechanism in which the cell population is divided into leader and follower states (McLennan *et al.*, 2012, 2015a), which are dynamically induced by microenvironmental signals (McLennan *et al.*, 2015b). In Xenopus cephalic NC, a complementary mechanism has been studied, based on balanced contact inhibition of locomotion and coattraction (Carmona-Fontaine *et al.*, 2011; Woods *et al.*, 2014), without explicit heterogeneity in the cell population.

Recent experiments in zebrafish used cell tracking evidence to argue for heterogeneity in trunk NC, but not cranial NC (Richardson *et al.*, 2016). Measures of heterogeneity used in that study include the directional correlation of cells with the migratory route, and change of relative position of cells within the group. As only a few tens of cells per embryo were analysed, an appropriate null model could help to answer how likely one is to see the differences observed in homogeneous populations. Furthermore, as cranial NC cells migrate in streams, and trunk NC cells in narrower chains, appropriate models could explore how different microenvironmental conditions with varying degrees of confinement can affect the chances of observing differences in measures of heterogeneity.

In our delay correlation analysis, we find that confinement tends to decrease the apparent heterogeneity (Fig. 3E), for a given parameter combination and population density. When including a fraction of 10% leader-like cells, confinement still decreases the measure of heterogeneity (compared to no confinement), but increasing confinement has less effect on the ability to detect heterogeneity (Fig. 3F). For similar analysis of in vivo tracking data from trunk and cranial NC cells, we can thus predict the following: If decreased heterogeneity was measured in the trunk vs head, the populations may, in fact, both be homogenous. This would suggest that leader-like cell states may not exist in vivo, as the observations (differences in heterogeneity) can be explained without them. If the measure of heterogeneity was greater in the trunk, both locations could host heterogeneous populations. Such an observation would support the notion that cells adopt leader-like states, or otherwise respond differentially to guidance cues, which could be of biological importance.

**Figure 3:**
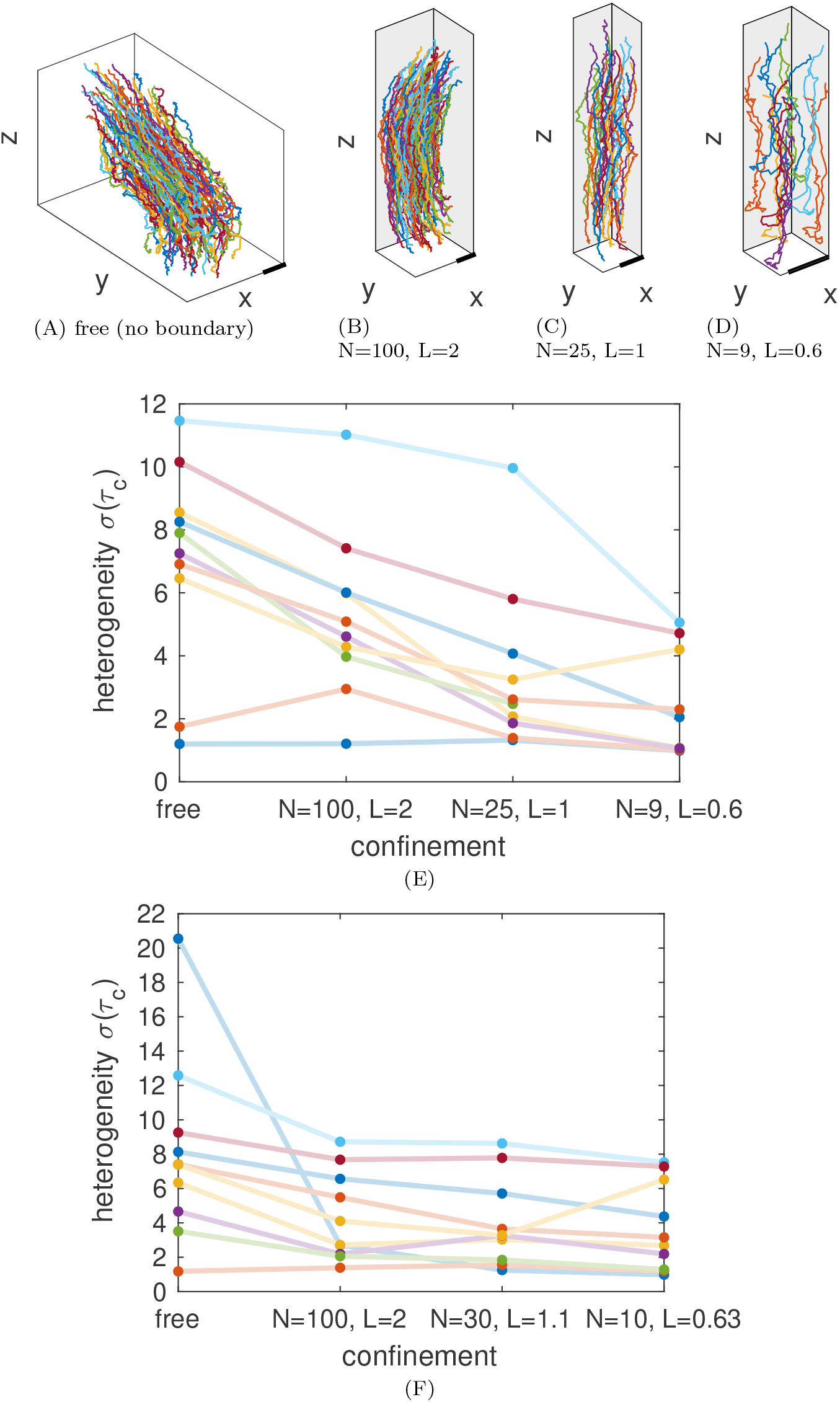
Heterogeneity under confinement. (A-D) Example trajectories of free (A) and confined (B: *L* = 2, C: *L* = 1, D: *L* = 0.6) simulations are shown for *α* = *β* = 4. No-flux boundaries are shown in grey, free boundaries in white, and the scale bar shows *L* = 0.5. Colours are chosen to distinguish different lines only. (E-F) The standard deviation of peak-delay times in directional cross-correlation, *σ*(*τ_C_*), for simulations with free boundary conditions as well as no-flux boundary conditions in *x*, *y* for three domain sizes, *L*. Cell number, *N*, was decreased to maintain the population density for different domain sizes. All parameter combinations in the integer range *α* ∈ {4, 8} and *β* ∈ {1, 2, 4, 8, 16} were simulated, with each line displaying results for a different parameter combination. Lines are to guide the eye only (horizontal axis is not continuous). (E) Results from homogenous cell populations. (F) Results with 10% of cells in a leader state (aligned with the *x*-axis). Note different scales in (E) and (F).

###### Tumor invasion

Heterogeneity of cell migration is of great interest to cancer research, for example to identify whether a subpopulation of cells in a tumor is more invasive, and how this invasive behaviour can be modulated by drugs or microenvironmental properties. Recent studies of mammalian cell cohorts in 3D environments have used delay correlation analysis to quantify heterogeneity (Sharma *et al.*, 2015). The authors found a range of delay distributions of different cell clusters, indicating potential evidence for heterogeneity of movement between different cell clusters (although making no direct claim as to any underlying heterogeneity of cell states themselves).

To distinguish between transient artefacts of small sample sizes and spatiotemporal heterogeneity as an intrinsic property of a cell collective, one has to compare the delay distributions with those generated from a suitable null model. To illustrate this, we simulated our model with lower cell numbers, representing the size of cell cohorts observed by Sharma *et al.* (2015) (see Comparison with experimental data). Our results (Fig. B1) show that existing data are broadly consistent with results obtained from a model without heterogeneity, as well as a model with a few cells striving to move in a particular direction. Here, heterogeneity cannot be inferred from the width of the peak-delay distribution alone. With heterogeneity, the model is best fit at lower interaction strengths for either dataset. The best fit for each of the datasets is at different simulation parameters, suggesting inter-cluster heterogeneity in the experimental observations, but a greater number of comparable experimental datasets is required to make any such inference robust. But the point of this illustration is explicitly not to argue that our model is the best description for the data, but that one cannot deduce heterogeneity from the measurements without consideration of an appropriate null model.

Furthermore, we found that stronger attraction-repulsion between cells can increase apparent heterogeneity in a population of identical cells (Fig. 2A). This leads to an intriguing hypothesis: When tracking cell clusters that undergo a transition to become more invasive, they may appear *less* heterogeneous as the cluster loosens and cells start to break free from the attachments to their neighbors.

## 3. Discussion

In this paper, we have used a minimal model of collective cell migration to highlight potential pitfalls in measuring heterogeneity from cell tracking data. We have presented a model for 3D collective cell migration that can generate trajectories of moving cell populations with a variety of collective behaviours (Fig. 1). Focusing on cohesively moving cell populations, we analysed the heterogeneity of migration using delay correlation analysis, demonstrating that non-zero heterogeneity can be measured in a homogeneous population with interactions (Fig. 2A). This apparent heterogeneity stemmed not just from cells with broad, but symmetric, individual delay distributions, but also from cells whose correlation with the rest of the population peaked consistently at non-zero delays (Fig. 2B), thus capturing leading and lagging on the timescales of observation. We further found that this bias towards apparent heterogeneity increases with stronger intercellular forces (Fig. 2C), but not consistently so with stronger alignment. By applying no-flux boundary conditions, we investigated how confinement affects the appearance of heterogeneity, and found that, for a given population density, more narrowly confined populations appear less heterogeneous (Fig. 3E). Finally, we suggested two biological applications where our results may be relevant: (A) in the study of neural crest cell migration in different embryonal microenvironments, and (B) when cancer cells undergo a transition to become more invasive (see Box: Biological applications). Based on the insight gained from our modelling study, we considered potential *in vitro* as well as *in vivo* experiments and hypothesized their outcomes.

Our goal was to present an illustrative example using a suitably generic model of collective movement, rather than to construct the most realistic model of collective cell migration – which will differ with each biological application. Even within this constraint, other modelling choices are possible: For example, the explicit alignment term in our model equations might seem unrealistic. In an alternative SPP model for collective cell migration, Szabó *et al.* (2006) have shown that short-range adhesive forces can be equivalent to an alignment term. We therefore anticipate that the results of our simulations and analysis would not change qualitatively if alignment of movement directions was mediated through another type of interaction, such as short-range adhesion.

To quantify heterogeneity of collective movement we chose the popular method of delay correlation analysis (Nagy *et al.*, 2010). The idea behind this method is to calculate to what extent some cells move first, and others follow. Alternative methods to determine the (directional) coupling between cells, other than cross-correlation, are available, such as causal information flow (Lord *et al.*, 2016; Richardson *et al.*, 2013), or “delay space” measures using the Fréchet distance (Konzack *et al.*, 2016). Other measures of heterogeneity (than delay in directional coupling) could be used to complement the analysis. One example is the degree of rearrangement of cells’ relative position, which can be quantified as “neighbor overlap” (Cavagna *et al.*, 2013).

Other studies have considered how to measure cell population heterogeneity in, for example, gene expression (Altschuler and Wu, 2010; Vallejos *et al.*, 2016). Our work is complementary as we show how even measuring distributions may not suffice to “determine which variation is random and which is meaningful” (Altschuler and Wu, 2010). Here, too, appropriate mathematical methods (Vallejos *et al.*, 2016), can be utilized to assess whether genetic heterogeneities of cell populations are statistically significant in the first place. Even if genetic heterogeneity can be reliably measured, it may still be of interest to correlate this with behavioural analysis, e.g., cell tracking.

*In vivo,* cell population heterogeneity is often located at the boundary of a migrating collective (McLennan *et al.*, 2015a), where cells are exposed to different microenvironmental signals (McLennan *et al.*, 2015b). Thus, it is natural to ask whether cells at the boundary of our simulated populations also show increased heterogeneity. Cells at the boundary have fewer neighbors to interact with than cells at the core, which influences the direction of movement through alignment, attraction-repulsion, and noise (2). However, when analysing cells at the boundary and core of the population separately, we found no difference in peak-delay distributions (Fig. S4). Therefore, the apparent heterogeneity that can arise from interactions between identical agents is not an ‘edge effect’.

In an extension of our model we included cells in a leader state, which align with a prescribed direction instead of with their neighbouring cells. For this particular form of cell population heterogeneity, different implementations could be considered within the SPP modeling framework. For example, Ferdinandy *et al.* (2017) adapted the model of Szabó *et al.* (2006) to include leader-follower heterogeneity by also varying the interaction strengths (albeit to represent horse harems, which also required directionality of interactions). Similarly, Chang *et al.* (2013) have used a SPP model with asymmetric alignment interactions to represent tumor-stromal interactions.

**Figure B1:**
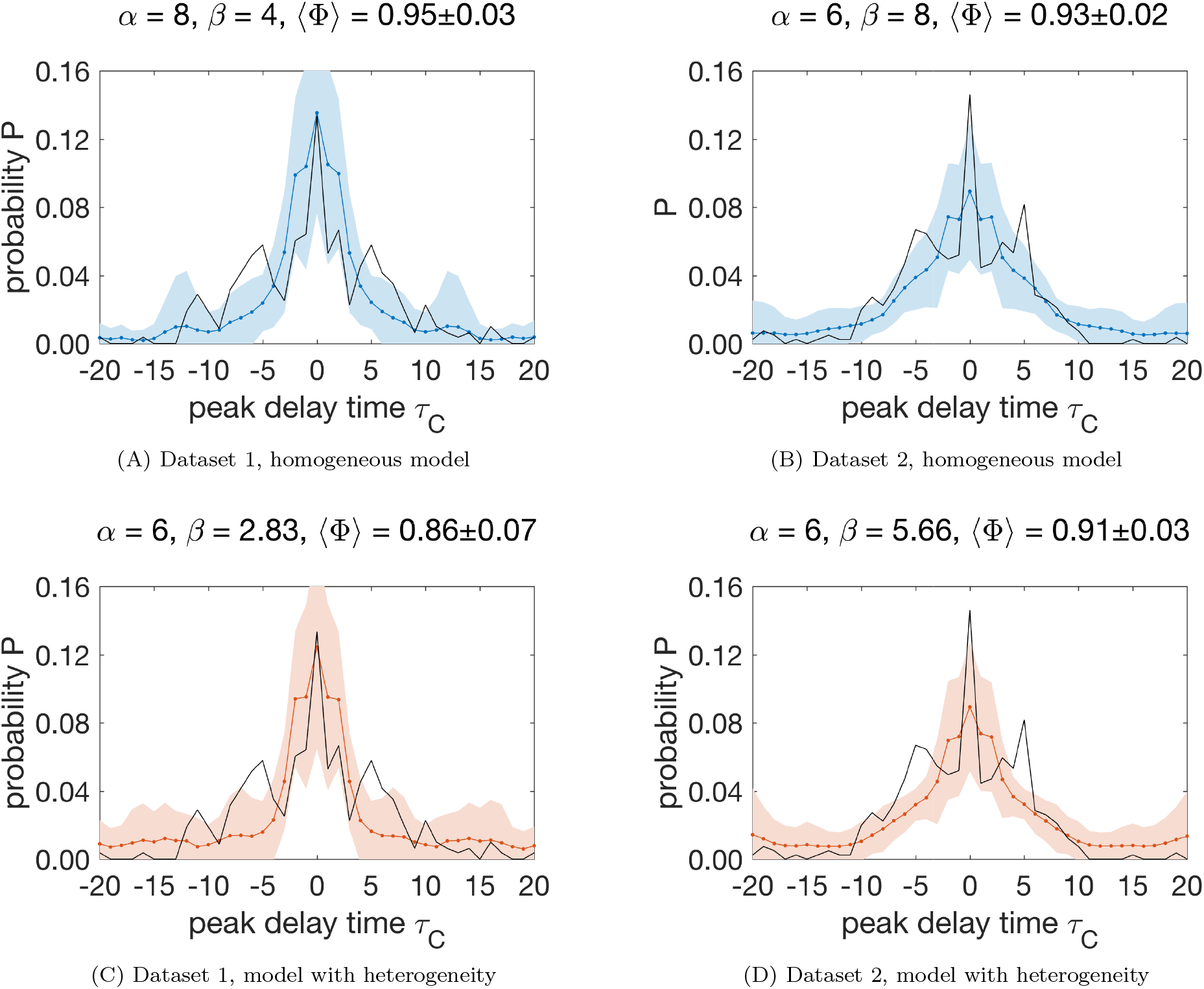
Comparison of models with and without heterogeneity with experimental data: (A) Experimental data (black line, estimated from Sharma *et al.*, 2015, Fig. 4C) and simulations for *N* = 10, *α* = 8, *β* = 4; (B) Experimental data (black line, estimated from Sharma *et al.*, 2015, Fig. 4D) and simulations for *N* = 20, *α* = 6, *β* = 8; (C) Experimental data (black line, estimated from Sharma *et al.*, 2015, Fig. 4C) and simulations for *N* = 10, *α* = 6, *β* = 2.83, with one informed cell; (D) Experimental data (black line, estimated from Sharma *et al.*, 2015, Fig. 4D) and simulations for *N* = 20, *α* = 6, *β* = 5.66, with two informed cells. Colored lines shows the mean of smoothed simulation results, shaded area shows 2*σ* confidence interval (*n* = 10). See main text, Comparison with experimental data, for details.

Our work illustrates how different choices of null models can affect the interpretation of heterogeneity in cell population data. Comparing different versions of our model, with and without heterogeneity, with experimental cell tracking data (Fig. B1), we showed that either was able to capture the width of the peak delay distribution, but different parameter values gave the best fit in each case. Without consideration of the homogeneous model, one may have been led to conclude heterogeneous motility in the tracked cell clusters. Even with both choices of null model to compare, one could not strongly differentiate, based on the data at hand, between a homogenous cell population and a less strongly interacting heterogeneous population – they yield peak delay distributions of similar width. This fact can be used by experimental researchers to assess limitations in their data, such as observation time, sampling frequency, and/or number of replicates.

Going beyond simulation studies, random matrix theory can be used to rigorously quantify the expected correlations between a collection of random variables. This branch of statistics has numerous and fruitful applications in physics, finance (Bouchaud and Potters, 2009), and, more recently, biology (Klein *et al.*, 2015). Analytical tractability of random matrix statistics, however, decreases drastically when venturing beyond independent Gaussian random variables. In collective cell migration, and in biology more generally, relevant null models fall into this territory more often than not. We may never see a random matrix theory of collective cell migration, but perhaps we can use it as inspiration to make headway with numerical calculation and computational modelling.

To close, we would like to loosely suggest best practices to avoid spurious correlations in complex biological systems. We ought to maximize observation time in a given experiment, as much as is reasonable without sacrificing the stability of the experimental system. When interpreting the results, it is imperative that we think carefully about what the null hypothesis is, and be aware that the usual tests for statistical significance may not apply. When comparing different biological systems, or observing the change in a system over time, we should also try to compare these with tailored, yet simple, simulations to sanity-check at least the qualitative differences or trends we observe. With this study, we hope to have contributed to a conceptual guideline for researchers interested in quantifying heterogeneity of cell populations, a field we expect to grow substantially with the increasing abundance of cell tracking data.

## Author contributions

Conceptualization, methodology, software, validation, formal analysis, investigation, data curation, writing – original draft, LJS; Writing – review & editing, visualization, supervision, project administration, LJS, PKM, and REB; Funding acquisition, LJS and REB.

## Acknowledgements

The authors thank Yasha Sharma, Diego Vargas, Muhammad Zaman, and Jeremy Green for insightful discussions. LJS was funded through a Doctoral Prize award by the UK Engineering and Physical Sciences Research Council (grant number EP/M508111/1).

## Supplemental information

### Method details

#### Intercellular forces

Intercellular forces point in the direction of the unit vector e_*ij*_ between cells *i* and *j*, i.e., **f**_*ij*_ = *f*(*r_ij_*)**e***_j_.* The magnitude of the force, *f*(*r_ij_*), was chosen to reflect hard core repulsion at short distances (informally described as a force of infinite magnitude), spring-like attraction-repulsion at intermediate distances, and no force at large distances. Specifically, the magnitude of the force between two cells *i* and *j* separated by a distance *r_ij_* is given by

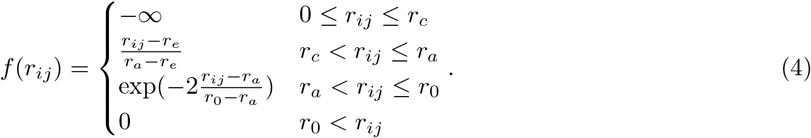

Here, *r_c_* denotes the core radius, *r_e_* the equilibrium cell separation, *r_a_* the attraction radius, and *r*_0_ the interaction radius, above which cells cease to exert forces on each other. Default parameter values used here are *r*_0_ = 0.2, *r_e_* = 0.5, *r_a_* = 0.8, and *r*_0_ = 1. In our numerical implementation we approximate the magnitude of infinite force (volume exclusion) by exp(100). The exponential regime was chosen to represent de-adhesion processes for increasing cell-cell separation. This does not accurately reflect forces for cells forming a new contact (decreasing distances), which would require a force-law exhibiting hysteresis. However, we have opted for the simpler form above as our simulations are of cell collectives in contact and we are here not investigating cases when these become cohesive after being initially separated. Thus, the form of intercellular force chosen is sufficient for our purpose. Figure S2 compares our choice of intercellular force with that of Grégoire *et al.* (2003).

### Quantification and statistical analysis

#### Global model behaviour

First, we quantify global alignment of velocities v*_i_* for *N* cells as

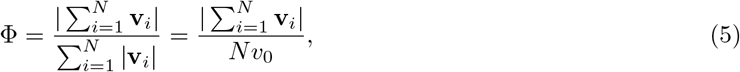

where the second equality holds for constant speed *υ*_0_. Thus, a value of Φ= 1 means all cells are perfectly aligned, while a value of Φ= 0 means complete disorder (or the special case of perfect anti-alignment, which we do not observe in our simulations). We use this order parameter to globally characterise the model behaviour, and thus restrict our analysis to instances of the model that correspond to the movement of ordered cell collectives only.

#### Comparison with experimental data

Sharma *et al.* (2015) conduct analysis of mammalian cell cohorts in 3D. In particular, they report broad peak-delay distributions of clusters of *N*_*c*1_ = 10 and *N*_*c*2_ = 19 cells. We chose to represent these with *N* = 10 and *N* = 20 cells in our computational experiments. To keep the cell density consistent, we chose the (random) initial cell positions to lie within a cube of length *L* = 0.9 and *L* = 1.1, respectively. Given these dimensions, our cell speed of *υ*_0_ = 0.05 approximately matches the displacement of cells (2.0–2.4 *μm* per frame, with an initial cluster diameter of ≈ 50*μm*, 2.4/50 = 0.048) seen in the datasets analysed. The experimental distributions were reproduced by reading the height of the bars of Figs. 4C&D in Sharma *et al.* (2015). To find matching peak-delay distributions, we simulated for a range of parameters and calculated the best fit as given by the root mean square deviation. The mean order parameter reported by Sharma *et al.* (2015) for these particular observations is 〈Φ〉_*c*1_ = 0.92 and 〈Φ〉_*c*2_ = 0.84. Hence, we restricted our comparison to simulation outcomes in which the order parameter was within 5% of this range.

**Figure S1:**
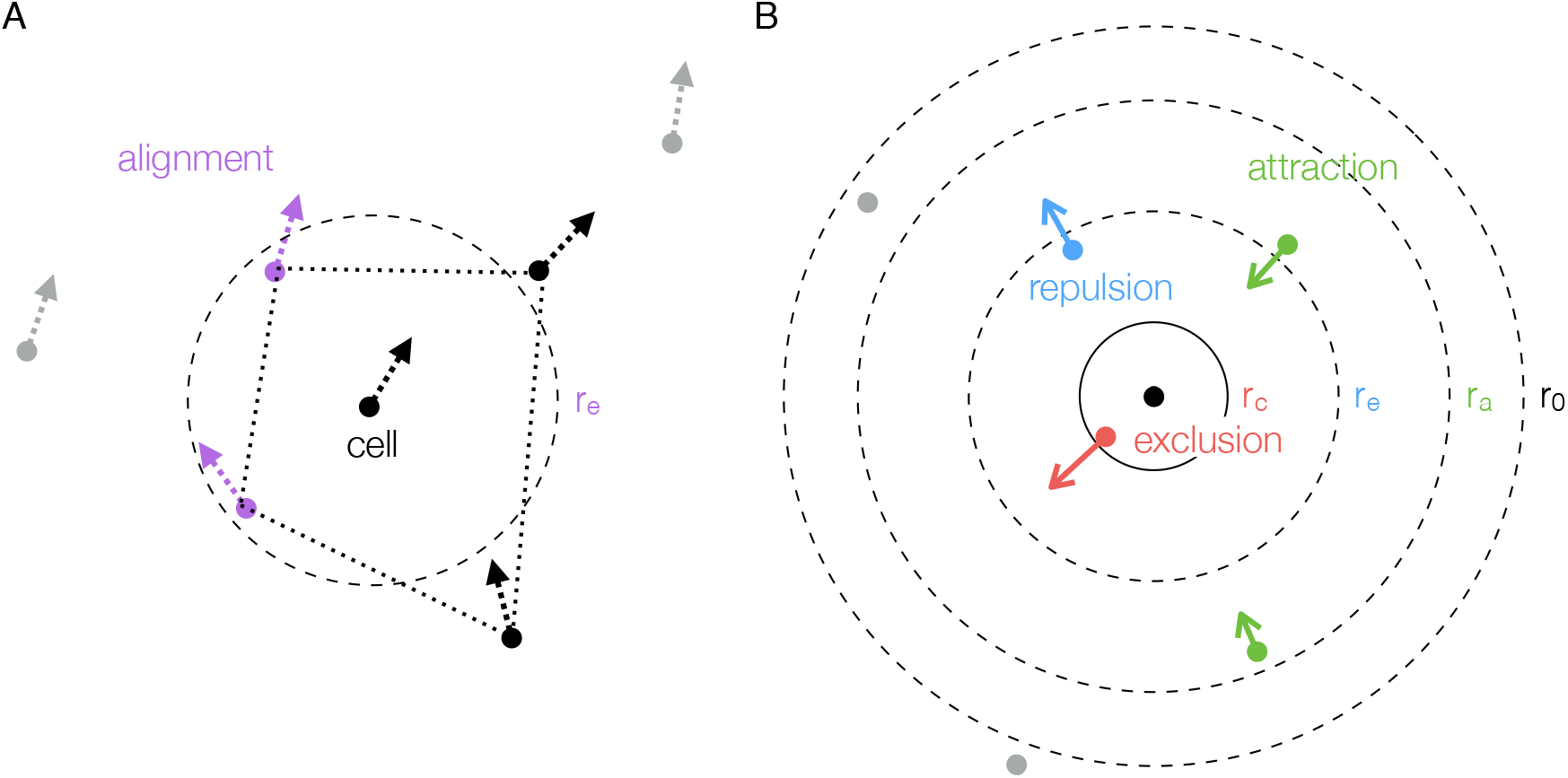
Schematic of interaction zones in the SPP model. Cells are drawn as dots with their direction of movement shown by dashed arrows (A), and the forces acting on them by solid arrows (B). (A) Alignment: A cell (drawn at the center of the circle) aligns with the direction of movement of neighboring cells within distance *r_e_* (shown in purple). Interactions are restricted to nearest neighbors, which are the vertices of the Voronoi region around the cell, as indicated by the dotted black line (here, cells within *r_e_* are a subset of the nearest neighbours, but this is not generally the case). Non-nearest neighbors shown in grey. (B) Intercellular forces: Cells are attracted or repelled from each other depending on their separation. Within *r_e_*, a repulsive force is exerted upon neighboring cells, shown in blue. At distances greater than *r_e_*, but smaller that *r*_0_, cells are attracted to each other, as shown in green. Thus *r_e_* is the preferred distance between cells. To model volume exclusion, cells exert a strong (approximately infinite) repulsive force upon neighbors within *r_c_*, shown in red. Non-nearest neighbors (shown in grey) do not interact, neither for alignment, nor for attraction-repulsion. Dashed lines mark the various interaction zones around the central cell. This schematic is drawn in two dimensions for simplicity, but applies analogously to the 3D model.

**Figure S2:**
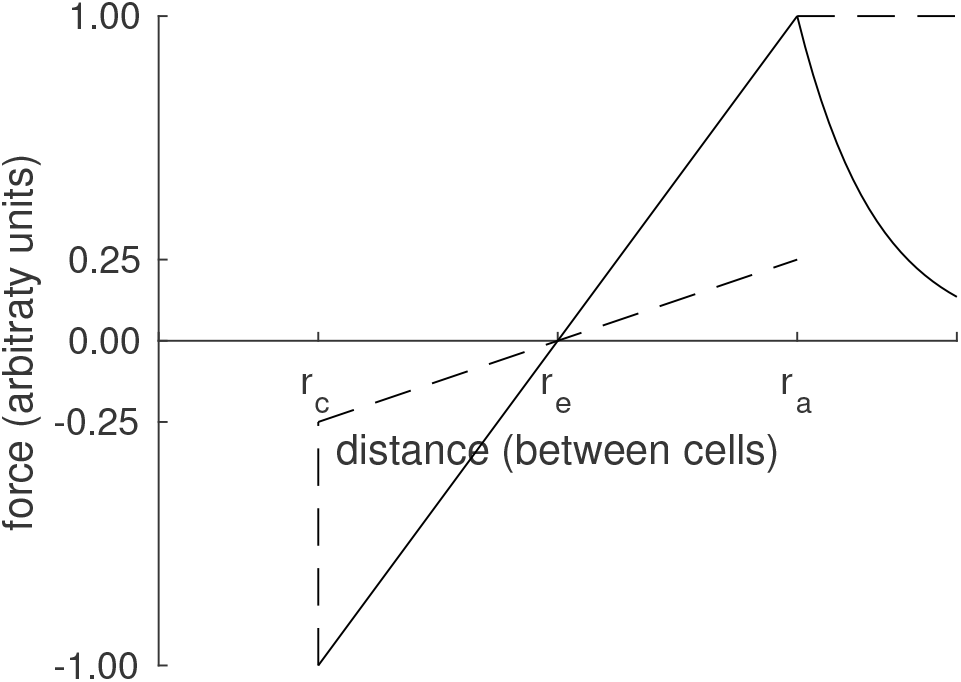
Intercellular forces, as in Grégoire *et al.* (2003) (dashed line), and the modified force used here (solid line). Positive forces are attractive, negative sign indicates repulsion. Below *r_c_*, the repulsive force is infinite, so that volume exclusion between cells is enforced (see main text for details).

**Figure S3:**
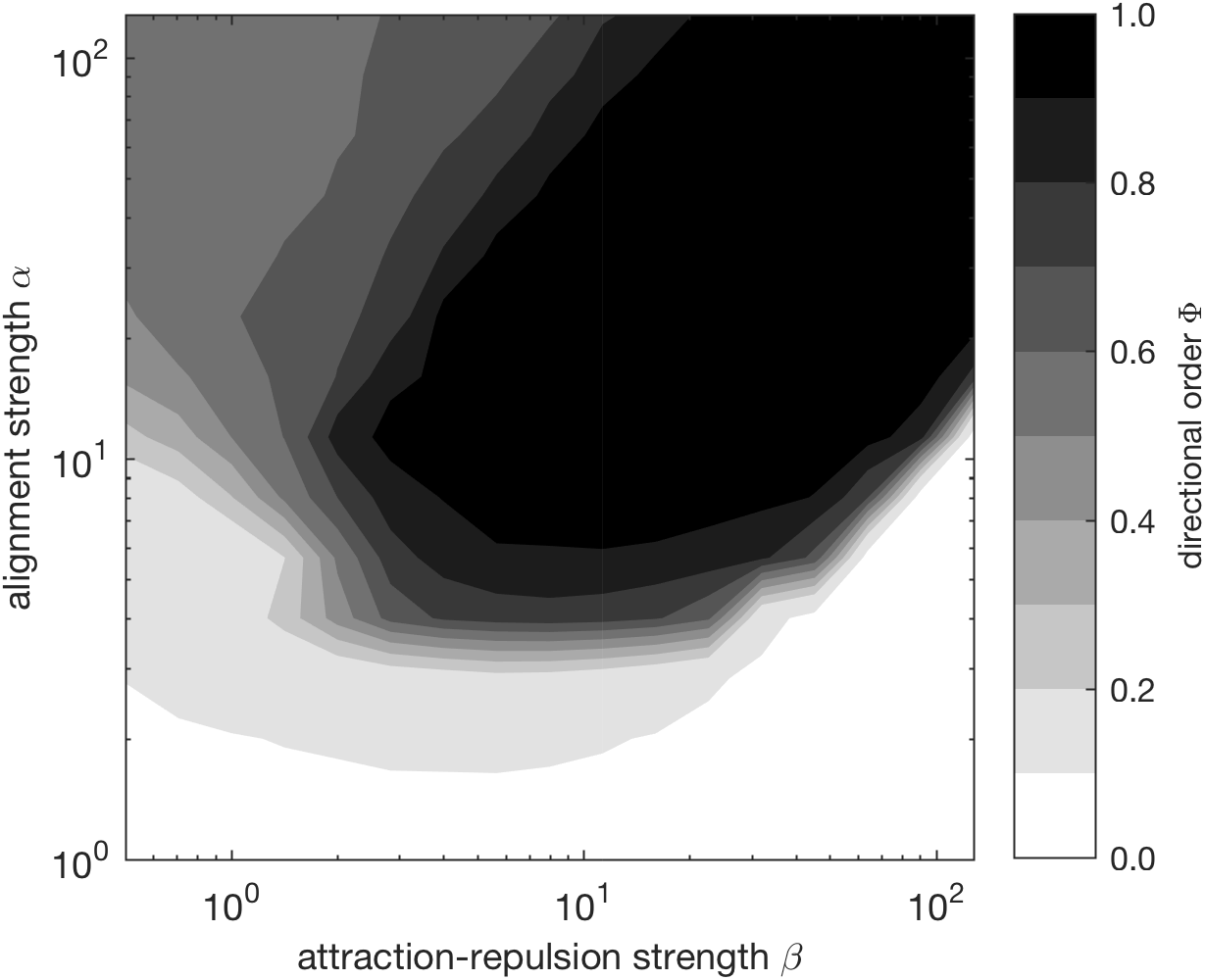
Global model behaviour, characterised by the phase diagram of the order parameter Φ, which is the overall alignment of cells (*N* = 100), averaged over 500 time-steps (after discarding the first 500 time-steps). Parameters were sampled on a log-spaced grid with 17 points in *β* and 15 points in *α*. For each parameter combination, the order parameter shown is an average over 10 simulations, with the same random number generator seed used for each run of every parameter combination.

**Figure S4:**
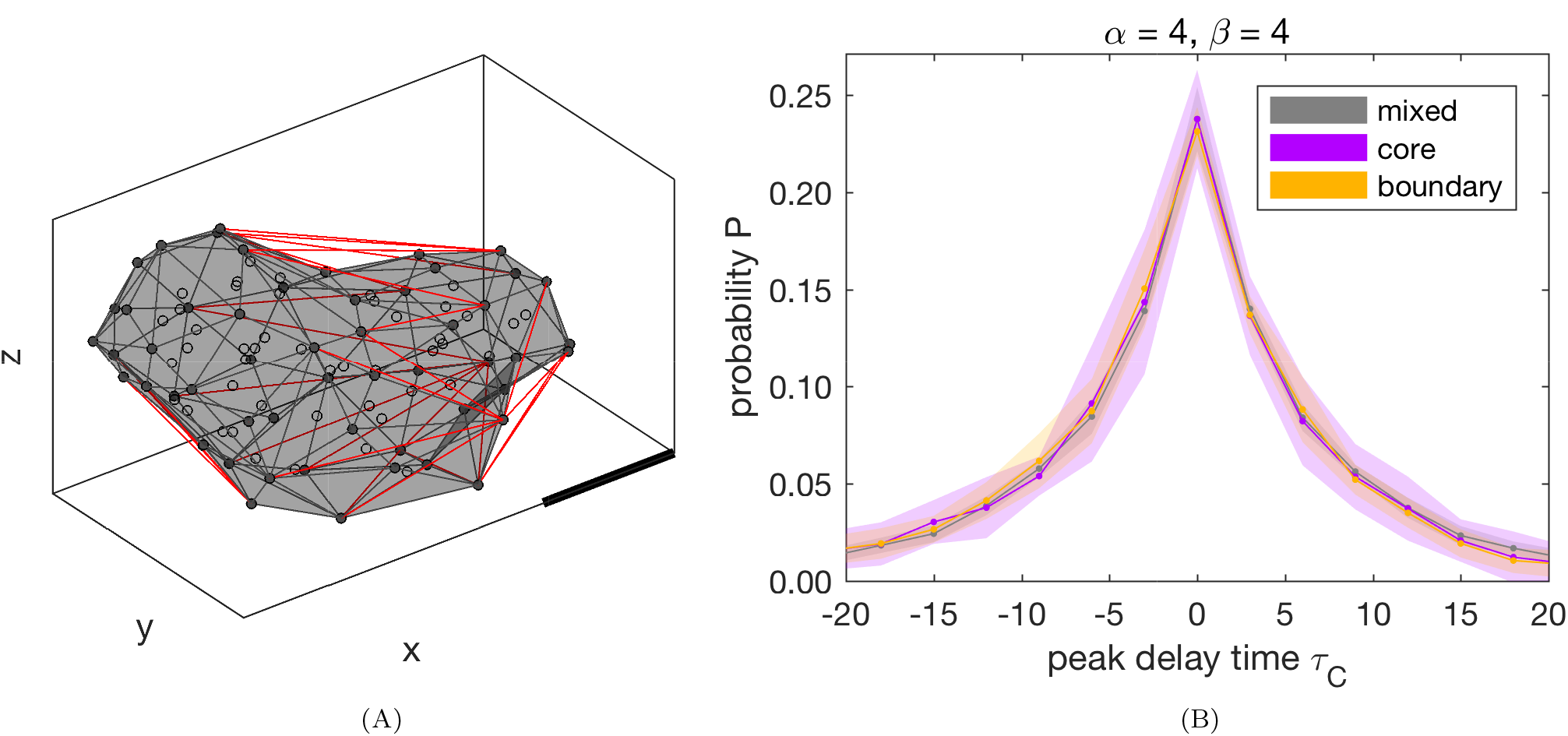
Apparent heterogeneity is not restricted to boundaries of the population. (A) Cells at the boundary are identified through a modified Delaunay triangulation: From a Delaunay triangulation of all cell positions (circles), we remove any edges (red lines) longer than the interaction distance, *r*_0_. The boundary of the remaining triangulation (grey surface) identifies the boundary cells (filled circles). (B) Peak-delay distributions for boundary, core, and mixed cells. As cells change their relative positions over time, we classify cells as ‘boundary’ or ‘core’ if they spend > 90% or % < 10% of time-points, respectively, at the boundary. Cells in neither category are classified as ‘mixed’. Points show the average for each bin over 10 simulations, shaded area shows the standard deviation.

